# Effect of *F. mosseae* on the expression of genes and proteins about sulfur nutrition in the continuous crop soybean roots

**DOI:** 10.1101/2022.10.19.512910

**Authors:** Xueqi Zhang, Chengcheng Lu, Ronglin Liu, Zixin Sun, Baiyan Cai

## Abstract

Soybean is a sulfur-loving oilseed crop, and continuous cropping can lead to soil sulfur deficiency, which can inhibit the growth and quality of soybean. This experiment used transcriptomic and proteomic sequencing techniques to analyse the changes in the expression of functional genes and related proteins in the root system of continuously cropped soybean and to reveal the molecular mechanism of *F. mosseae* inoculation on the soybean root system in response to sulfur nutrient supply at the molecular level. It was thus demonstrated that *F. mosseae* could enhance the uptake and transport of soil sulfur in continuously cropped soybean. This study, therefore, provides a theoretical basis for the application of *F. mosseae* as a biofertilizer in soybean production on sulfur-deficient soils.

**One-sentence summary:** *F. mosseae* affects soybean genes and proteins at the transcriptome and proteome levels.

## INTRODUCTION

Sulfur(S) is one of the essential nutrients for crop growth; is a basic building block for sulfur-containing amino acids as well as some enzymes, thioredoxin and other essential compounds (Iciek et al., 2019); and has important physiological functions in crop growth, development and metabolism. Sulfur deficiency can lead to yellowing of young plant leaves, reduced biomass, elongated stem tissue, interveinal necrosis and slow growth, ultimately leading to a significant reduction in yield and quality (Lunde et al., 2010; Farooq et al., 1999; Wali et al., 2004). Sulfur is usually taken up in the plant root system as sulfate, and small amounts of volatile sulfur (e.g., H_2_ S, SO_2_ and dimethyl sulfate (DMS)) can be taken up and metabolized by plant leaves (Mark et al., 2004). Sulfur also plays an important role in the construction of plant photosynthetic organs as well as in the electron transport system, and the reduced photosynthetic capacity of plants due to sulfur deficiency has been observed in plants such as maize (Antal et al., 2018; Vally et al., 2022). It has also been shown that sulfur application increases the sulfur and cysteine content of soybean leaves and seed tissues. Sulfate metabolism in higher plants occurs mainly in plastids, especially in chloroplasts (Liu et al., 2019). The assimilation of sulfate in plants can be divided into sulfate uptake and transport, sulfate activation, sulfate reduction and Cys synthesis. Cys synthesis is the only way for reduced sulfur to be metabolised, and the reduction of sulfide from SO_4_^2−^to S_4_^2−^occurs only in the plastids of plant cells. Sulfur from the soil and atmosphere is mainly taken up by plants as SO_4_^2−^ (Singh et al., 2018) and is subsequently reduced and assimilated by enzymatic catalysis to form cysteine (Cys) in the organic carbon skeleton. Plants use Cys as a precursor to synthesize a range of metabolites (e.g., glutathione) that are essential for growth and development (Takahashi et al., 2010).

Soybean (*Glycine max* (L.) Merr.) is a sulfur-loving crop, and its nutritional value is limited by its sulfur-containing amino acid content. The critical concentration of sulfur deficiency and the toxic concentration of sulfur in soybean are 0.194% and 0.277% (dry weight), respectively. Sulfur deficiency and toxicity have an inhibitory effect on soybean seed yield, and the content of seed storage proteins (albumin, globulin, glutenin and alcohol-soluble glutenin) is also reduced under sulfur stress conditions (Chandra et al., 2016). Increasing the efficiency of sulfate assimilation and utilization is an effective way to increase the sulfur-containing amino acid content of soybean seeds (Koralewska et al., 2008). A variety of sulfate transporter (SULTR) proteins act in concert during the uptake of sulfur by the plant body. These sulfate transporter proteins are present in different tissues, are not expressed at the same level and have different affinities. The high-affinity sulfate transporter is expressed specifically in the roots and is responsible for the uptake of SO_4_^2−^from the soil, whereas the low-affinity sulfate transporter is mainly expressed in the leaves and is responsible for the transport of sulfur in the plant. The soybean sulfur transporter gene *GmSULTR1;2b* is specifically expressed in roots and is induced in response to sulfur deficiency (Maruyamanakashita et al., 2004). Sulfur deficiencies are becoming increasingly pronounced in agricultural fields. The reasons for this are, on the one hand, the decrease in the number of organic fertilizers applied to farmland, the gradual decrease in soil organic matter, and the gradual replacement of sulfur-containing fertilizers (ammonium sulfate, potassium sulfate mixed with calcium superphosphate, etc.) by high concentrations of sulfur-free fertilizers (urea, diammonium phosphate) and the gradual decrease in the return of sulfur to farmland soils (Buchner et al., 2010). On the other hand, it is the progressive increase in sulfur taken away from the soil by crops due to increasing crop yields and increasing replanting indices. Continuous cropping results in selective uptake of nutrients from the soil (Yue et al., 2020).

Arbuscular mycorrhizal fungi (AMF) are beneficial soil microorganisms that can form a symbiotic relationship with the nutritional root systems of more than 80% of terrestrial plants, a relationship that is commonly found in nature (He et al., 2017). Mycorrhizal fungi not only use their symbiotic relationship to promote the uptake of N, P, K, minerals and water by plants but also perform well in assisting plants in increasing their resistance to stress and disease (Davison et al., 2015; Gao et al., 2019; Gomez-Bellot et al., 2014; Sun et al., 2019; EiFaiz et al., 2015; Ma et al., 2019; Ortiz et al., 2015). On the other hand, mycorrhizal fungi play a very important role in ecosystems, such as improving the physicochemical properties of plant inter-root soils, regulating the secretion of secondary metabolites by plant roots, and activating and inducing host defence systems and other related mechanisms (Gao et al., 2017; Mayer et al., 2017; Hakeem et al., 2016; Garg et al., 2010; Zhang et al., 2014). It has been shown that mycorrhizal plants are less affected in S-deficient soil habitats (Wipf et al., 2014), which is related to the presence of AMF, and therefore, it is believed that AMF have great potential value in improving crop yield in sulfur-deficient areas (Gahan et al., 2014). Sulfur affects the diversity, abundance values and colony structure of AMF in continuous soybean soils; sulfur has an impact on AMF diversity, abundance values and flora structure in continuous crop soybean soils; and appropriate sulfur levels can increase soybean yield and improve soybean quality (Bai et al., 2013; Li et al., 2012; Wei et al., 2013; Yu et al., 2013). However, how AMF affects the regulation of sulfur nutrient uptake in soybean and which proteins and enzymes are involved have not been addressed or clarified.

To address these questions, this study used proteomic sequencing and transcriptomic sequencing techniques to identify differentially expressed genes (DEGs) and differentially expressed proteins (DEPs) in soybean roots during the peak sulfur uptake period of continuous soybean crops based on the previous studies mentioned above. This study was conducted to identify key genes that influence the sulfur metabolism process in soybean and to reveal the effect of *F. mosseae* inoculation on sulfur uptake and transport in soybean roots at the gene and protein levels; investigate the changes in the content of genes and proteins related to sulfur uptake, translocation and assimilation in soybean roots; elucidate the mechanism of differential expression of sulfur-related proteins and genes; and provide a theoretical and practical basis for the application of AMF fungicide as a biological agent for nutrient remediation in continuous cropping soils.

## RESULTS

### AMF colonization, physical growth parameters and root activity

AMF colonization was analysed by taking the inoculated FS group as an example (Figure 1a). At the beginning of colonization, a small amount of mycelia was produced; at the middle stage, mycelia density increased significantly, and intra-root spores were observed. By the end, mycelia infested almost all fibrous root systems (≧95%). The results were obtained by calculating the colonization rate (CR). At 75 d, the maximum colonization rate was reached. The increase in the colonization rate of the S-added treatment group (NS, FS) slowed down after 30 d of sulfur treatment, probably due to the preference of AMF itself, which to a certain extent weakens AMF infestation in the short term when the environmental nutrient conditions are good, while a moderate supply of exogenous sulfur can increase the intensity of mycorrhizal infestation. The colonization rates of FS and NS were consistently higher than that of N. The colonization rate of soybean roots grown in continuous cropping soil (N) was significantly lower than that of soybean roots grown in uncropped soil (CK), indicating that continuous cropping reduces the colonization rate of plants by beneficial fungi in the soil (Figure 1b).

**Figure 1.**
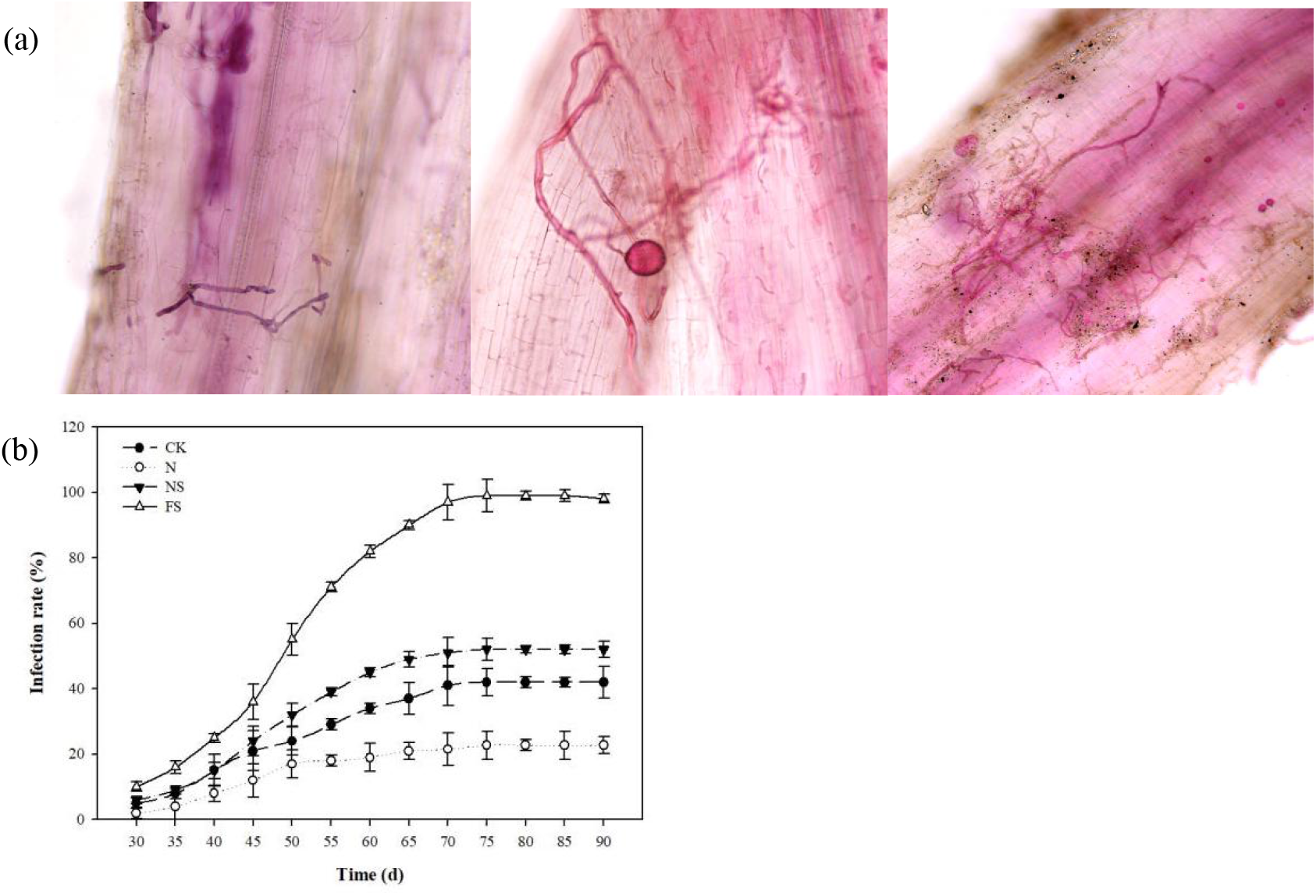
AMF colonization, physical growth parameters and root activity. (a) AMF colonization status of different growth stages in FS; (b) Arbuscular mycorrhizal infection rate. CK, control group, HN48 was sown in first-year soybean soil; N, HN48 was sown in three-year-old soybean soil; NS, HN48 was sown in three-year-old soybean soil with 0.5 mmol/L MgSO_4_; FS, HN48 was sown in three-year-old soy*bean soil* inoculated with *F. mosseae* and 0.5 mmol/L MgSO_4_. The abscissa represents the days after sowing.

Under the effect of the inoculation treatment, plant height varied significantly between groups at 60 d after sowing, with the FS group heights gradually approaching those of the CK group. Plant height grew slowly between 60 d and 75 d for the CK and FS groups and was almost equal by 75 d. The dry and fresh weight of soybean plants increased significantly between 30 d and 60 d. After 60 d, the change in dry and fresh weight of soybean plants levelled off (Figure S1). Based on observations during the pod set, the number of pods in the NS and FS groups was higher than that in the CK and N groups. The addition of S caused the soybean plants to use more nutrients for seed maturation later in the growing season. A moderate supply of exogenous sulfur and inoculation with *F. mosseae* increased soybean plant height.

Root activity correlates with planting survival and is a key indicator of seedling quality. In both the inoculated and control groups, root activity peaked at 60 d, when soybean root growth activity was high and vigour increased. The effect of the continuous crop on root activity was also significant, with root activity in the uncropped CK group significantly higher than in the uncropped N treatment group. The root system then began to senesce between 75 and 90 d, and vigour declined (Figure 2).

**Figure 2.**
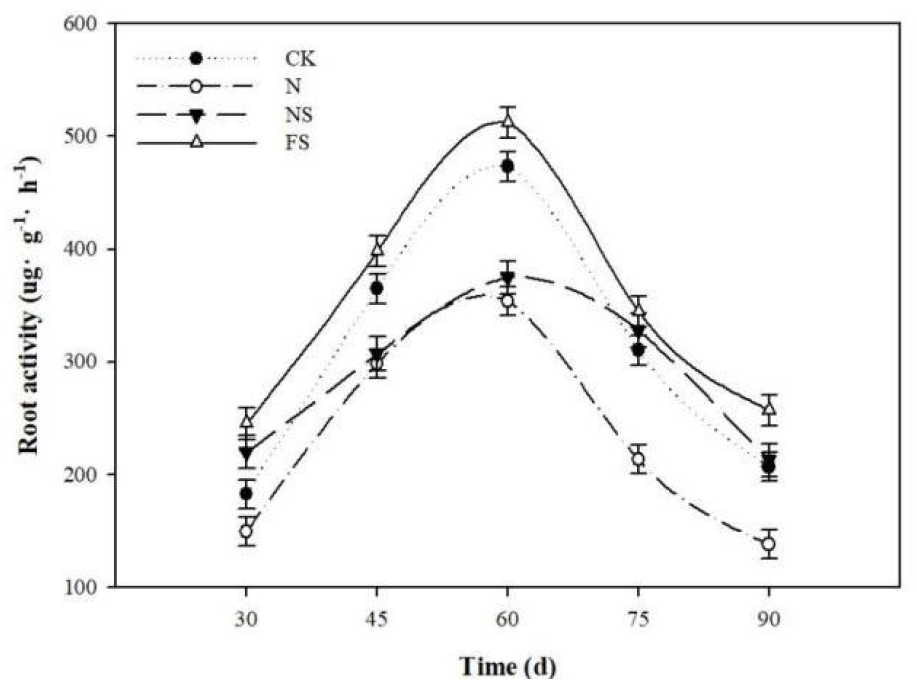
Root activity of soybean plants. CK, control group, HN48 was sown in first-year soybean soil; N, HN48 was sown in three-year-old soybean soil; NS, HN48 was sown in three-year-old soybean soil with 0.5 mmol/L MgSO4; FS, HN48 was sown in three-year-old soybean soil inoculated with F. mosseae and 0.5 mmol/L MgSO4. The abscissa represents the days after sowing.

### Effects on sulfate uptake into roots, transportation and assimilation

The total sulfate content in both the functional leaves and roots of soybean showed a trend of increasing and then decreasing as fertility progressed. The total sulfate content in functional leaves was highest in the CK, N and NS groups without AMF at 45 d. The total sulfate content in the root system was highest in the CK, N, NS and FS groups at 60 d. In the FS group, it was highest at 60 d when the mycorrhizal infestation was more active. The total sulfate content of soybean leaves and roots was significantly influenced by exogenous sulfur and the sulfate concentration, as shown by the results of total sulfate content measurements. During the nutritional growth of the plant, most of the sulfate absorbed by the root system goes to the developing leaves, as these are the main sites of protein synthesis (Figure 3). Inoculation with *F. mosseae* significantly increased the uptake of soil sulfur in soybean and was most effective in promoting sulfate uptake during periods of active mycorrhizal infestation. During reproductive growth, sulfate is supplied mainly to the seeds, and sulfate is transported from the roots and leaves to the seeds, leading to a steady decline in sulfate content in the later stages of growth. Sulfate is not only a component of protein amino acids but is also involved in regulating the formation of disulfide bonds in proteins and enzymes (Om et al., 2022).

**Figure 3.**
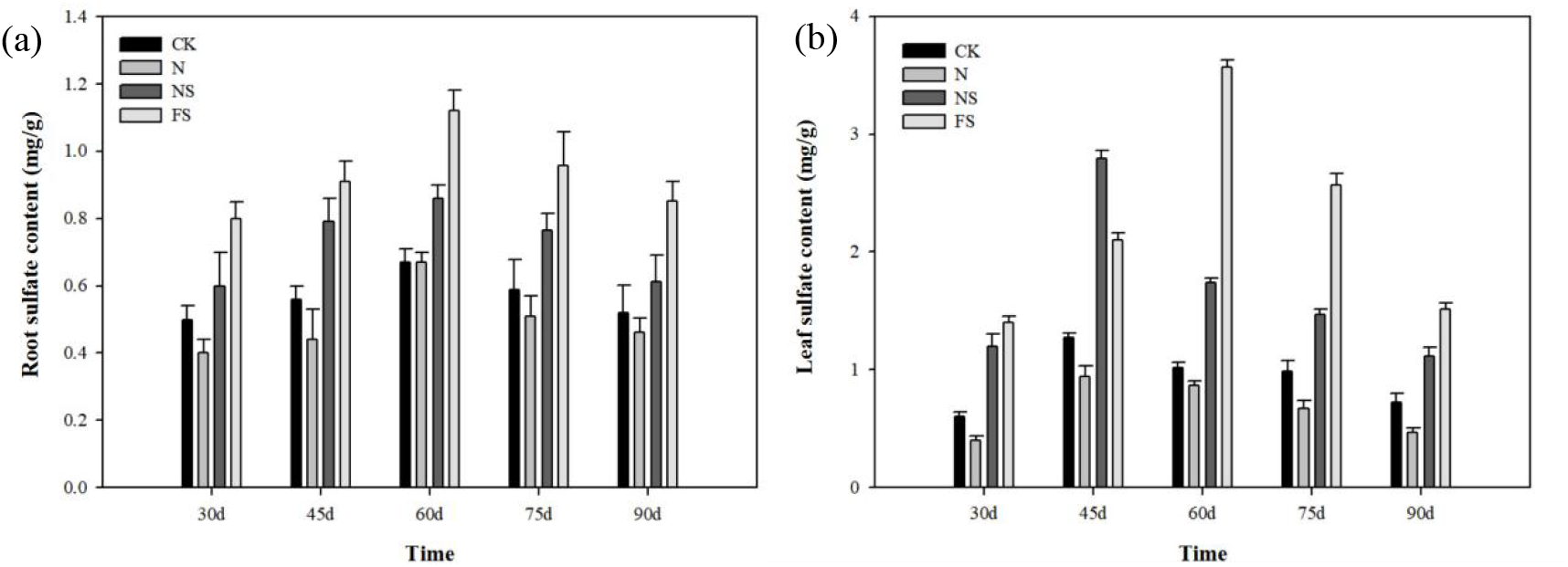
Sulfate content of soybean plants. (a) Leaf sulfate content.; (b) root sulfate content. CK, control group, HN48 was sown in first-year soybean soil; N, HN48 was sown in three-year-old soybean soil; NS, HN48 was sown in three-year-old soybean soil with 0.5 mmol/L MgSO4; FS, HN48 was sown in three-year-old soybean soil inoculated with F. mosseae and 0.5 mmol/L MgSO4. The abscissa represents the days after sowing.

The results of measuring ATPS and cysteine synthase activities in soybean roots showed that the ATPS enzyme activities of the roots of the CK and N groups showed a trend of first increasing and then decreasing, and it was significantly higher in the continuous crop group N than in the noncontinuous crop group CK. The enzyme activities of the NS and FS group showed a trend of first decreasing and then increasing from 30 to 60 days, and the enzyme activities decreased from 60 to 90 days (Figure 4).

**Figure 4.**
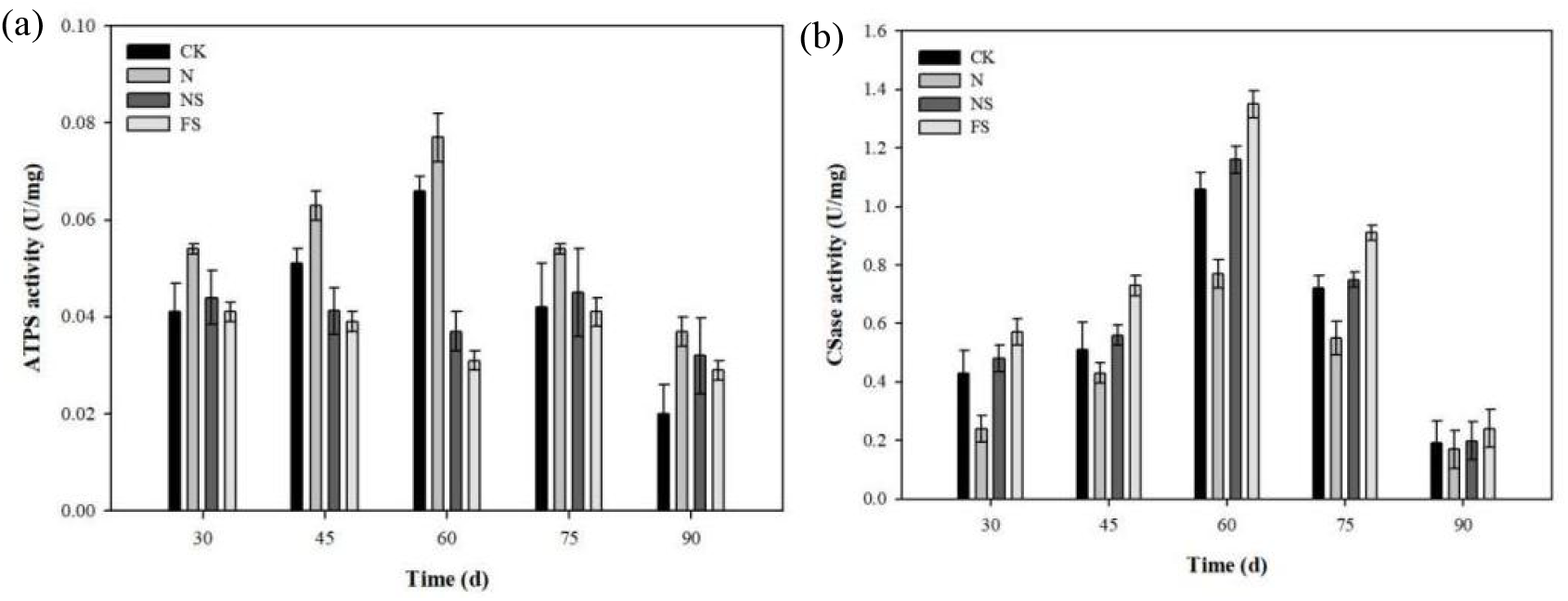
The activity of key enzymes in sulfur metabolism. (a) ATPS activity; (b) CSase activity. CK, control group, HN48 was sown in first-year soybean soil; N, HN48 was sown in three-year-old soybean soil; NS, HN48 was sown in three-year-old soybean soil with 0.5 mmol/L MgSO4; FS, HN48 was sownin three-year-old soybean soil inoculated with F. mosseae and 0.5 mmol/L MgSO4. The abscissa represents the days after sowing.

Soybean is an important source of plant protein (Li et al., 2017). The results of soluble protein content measurements in the roots and leaves of soybean plants showed that the CK and N groups reached the highest levels of protein content in the roots and leaves at 45 d, with decreasing levels in the later stages of flowering, and the NS and FS groups reached the highest levels in the roots and leaves at 75 d (Figure S2).

### Transcriptomic Response in Soybean Plants Inoculated with AMF

Significantly different genes were identified based on the criteria of FDR<0.05 and |log2FC|>1. Four comparative combinations were set up for two-by-two comparisons: CK-vs.-N,N-vs.-NS, NS-vs.-FS and CK-vs.-FS. In CK-vs.-N, a total of 49 DEGs (DEGs) were identified, of which 28 were upregulated and 21 were downregulated. In N-vs.-NS, a total of 23 DEGs were identified, of which 4 were upregulated and 19 were downregulated. In NS vs. FS, a total of 1906 DEGs were identified, of which 900 were upregulated and 1006 were downregulated. In CK-vs.-FS, only 27 DEGs were identified, of which 15 were upregulated and 12 were downregulated. To reflect the effect of sulfur addition versus inoculation with *F. mosseae* on gene expression trends, a trend analysis was performed for CK vs. N vs. NS vs. FS. DEGs were co-classified into 20 gene expression patterns, including seven upregulated expression patterns in NS (including profile 11, profile 4, profile 5, profile 12, profile 18, profile 6, and profile 19) and 7 upregulated expression patterns in FS (including profile 8, profile 3, profile 10, profile 15, profile 1, profile 6, and profile 19) (Figure S3).

GO analysis was performed to classify the functions of the annotated unigenes. These unigenes were categorized into three main GO categories; after inoculation treatment (NS vs. FS), in the biological process category, metabolic process (618, 32.4%), cellular process (519, 27.2%) and single-organism process (380, 19.9%) were dominant. In the component category, membrane (145, 7.6%), cell (106, 5.6%) and cell part (106, 5.6%) were the most highly represented. The most common terms were binding (597; 31.3%), catalytic activity (592; 31.1%) and nucleic acid binding transcription factor activity (100; 5.2%) (Figure 5a).

**Figure 5.**
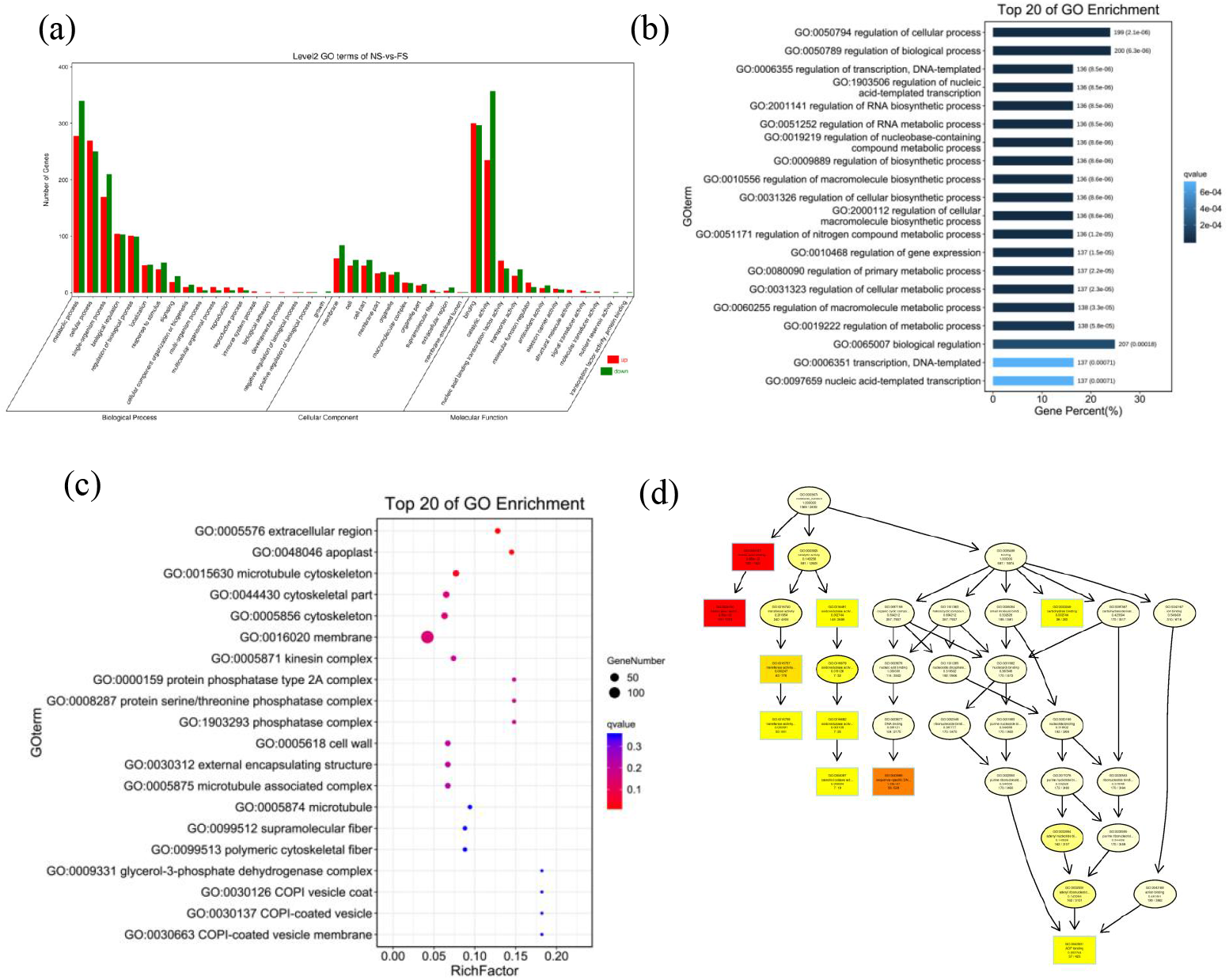
The result of GO enrichment analysis in NS vs. FS DEGs. (a) Go enrichment classification bar chart. The abscissa is the level 2 Go term and the ordinate is the number of genes in the term. Red indicates upregulation and green indicates downregulation. (b) Bar chart of biological process category. The top 20 GO terms with the smallest Q value are used for mapping. The top 20 Go terms with the smallest Q value are used for mapping. The ordinate is GO terms, and the abscissa is the percentage of the number of Go terms in the number of all differential genes. The darker the color, the smaller the Q value. The value on the column is the number and Q value of the Go terms. (c) Bubble diagram of cellular component category. The ordinate is Go terms, and the abscissa is enrichment factor (the difference genes in the Go terms are divided by all numbers). The size indicates the number. The more red the color, the smaller the Q value. (d) Interaction network graph of molecular function category.

In the biological process category, the top 20 GO terms (Q value < 0.05) are mostly associated with “regulation” such as regulation of cellular process; regulation of biological process; regulation of transcription, DNA-templated; regulation of nucleic acid-templated cellular component enriches some of the GOs, such as regulation of cellular process; regulation of biological process; regulation of transcription, DNA-templated; regulation of nucleic acid-templated transcription; and regulation of RNA biosynthetic process (Figure 5b). In the cellular component category, some GO terms were enriched, which are related to the extracellular region, apoplast, and microtubule cytoskeleton (Figure 5c). The majority of DEGs were involved in the molecular function category and related to GO terms such as nucleic acid binding transcription factor activity, transcription factor activity, sequence-specific DNA binding, catechol oxidase activity, oxidoreductase activity, and carbohydrate. GO terms such as nucleic acid binding transcription factor activity, transcription factor activity, sequence-specific DNA binding, catechol oxidase activity, oxidoreductase activity, carbohydrate binding, and ADP binding were significantly enriched (Figure 5d).

### Results of annotation and enrichment analysis of the DEGs KEGG pathway

The pathway-based analysis can help to further understand the function of different genes in coordinating their biological behaviour, so KEGG was used to annotate the information of the identified genes at the pathway level. The top 20 enriched pathways in NS vs. FS included secondary metabolite biosynthesis, phenylpropanoid biosynthesis, plant-pathogen interaction, MAPK signalling pathway-plant, linoleic acid metabolism, cyanoamino acid metabolism, isoquinoline alkaloid biosynthesis, alpha-linolenic acid metabolism, etc. (Figure 6). In NS-vs.-FS, 67 DEGs were annotated to 8 pathways (phenylpropanoid biosynthesis. Isoquinoline alkaloid biosynthesis, flavonoid biosynthesis, glucosinolate biosynthesis, etc.), these pathways belong to the same biosynthesis of other secondary metabolites; 69 DEGs were annotated to 3 pathways (MAPK signalling pathway - plant, plant hormone signal transduction, phosphatidylinositol signalling system), which are part of the signal transduction; and 78 DEGs were annotated to 13 pathways (cysteine synthase and methionine metabolism, valine, leucine and isoleucine biosynthesis, tyrosine metabolism, and glycine, serine and threonine metabolism) (Table S1). In N-vs.-NS, cysteine synthase and methionine metabolism are associated with amino acid metabolism. The methionine metabolism and sulfur metabolism pathways were both annotated to show significant downregulation of cysteine synthase (*cysK*) expression. In NS-vs.-FS, the cysteine and methionine metabolism pathways annotated serine O-acetyltransferase (*cysE)*, cysteine synthase (*cysK*), and O-acetylhomoserine/O-acetyl serine sulfhydrylase (*MET17)* as significantly upregulated. The sulfur metabolism pathway showed significant upregulation of serine O-acetyltransferase (*cysE)*, cysteine synthase (*cysK, cysO, ATCYSC1)*, and o-phosphatidylserine sulfhydrylase/cystathionine beta synthase (*ATCYSC1*), while 3’(2’), 5’-diphosphate nucleotidase (*cysQ, MET22, BPNT1*), Golgi PAP phosphatase (*IMPAD1, IMPA3*) and other phosphatase genes were significantly downregulated (Table S2).

**Figure 6.** The result of KEGG enrichment analysis in NS vs. FS DEGs. KO enrichment bubble diagram. The top 20 pathways with the smallest Q value are used for mapping. The ordinate is pathway, and the abscissa is enrichment factor (the number of differential genes in the pathway is divided by all the numbers). The size indicates the number. The more red the color, the smaller the Q value).

### GSEA (Gene Set Enrichment Analysis) of DEGs

To avoid missing some important DEGs affected by inoculation with AMF, the author performed GSEA -GO and GSEA-KEGG analysis on nonsignificantly enriched differential genes and showed 20 GO terms each for the NS-vs.-FS group NES maximum and minimum.

The GSEA-GO results showed in NS vs. FS, GO terms related to plant cytoskeleton: cytoskeleton, microtubule cytoskeleton, cytoskeleton protein binding; GO terms related to membrane transport: amino acid transmembrane transporter; GO terms related to membrane transport: amino acid transmembrane transporter activity, organic acid transmembrane transporter protein activity, organic anion transmembrane transporter protein activity, and phosphatase complex; GO terms related to ARF (auxin response factor): regulation of ARF protein signal transduction, ARF protein signal transduction; GO terms related to ARF (auxin response factor): regulation of ARF protein signal transduction, ARF protein signal transduction, and ARF guanylate exchange factor activity; GO terms related to plant signal transduction: regulation of intracellular signal transduction, Ras protein signal transduction, and regulation of small GTPase-mediated signal transduction were significantly enriched (Table S3).

The GSEA-KEGG results showed that in NS vs. FS, enriched pathways related to plant defence responses included autophagy-other eukaryotes, the plant MAPK signalling pathway, flavonoid and flavanol biosynthesis, flavonoid biosynthesis, tropane, piperidine and pyridine alkaloid biosynthesis; pathways related to fatty acid biosynthesis, fatty acid degradation, and fatty acid metabolism; amino acid anabolic pathways included beta-alanine metabolism, valine, leucine and isoleucine degradation, and cyanogenic amino acid metabolism; and plant terpenoid synthesis pathways included terpenoid backbone biosynthesis, diterpenoid biosynthesis, sesquiterpene and triterpene biosynthesis (Table S4).).

### qRT-PCR Confirmation of Selected Genes Expression Levels

To verify the reliability of the RNA-seq data, five DEGs were randomly selected for qRT - PCR analysis for transcript-level expression (Figure S4). The expression trends of the differentially expressed transcripts in the soybean roots during the period of the highest infestation rate were consistent with the sequencing results, indicating that the sequencing results were accurate and reliable.

### Proteomic Response in Soybean Plants Inoculated with AMF

To understand the underlying mechanisms of the physiological response to AMF inoculation, proteomic analysis was performed on the root samples harvested 75 d after sowing.

The DEPs were subjected to GO annotation analysis. The top 20 GO terms in NS vs. FS show that the main molecular functions exercised by DEPs include peroxidase activity, antioxidant activity, oxidoreductase activity, damaged DNA binding, hydrogen ion transmembrane transporter activity, monovalent inorganic cation transmembrane transporter activity, cation transmembrane transporter activity, and inorganic cation transmembrane transporter activity. The major components of cellular components include the intrinsic component of the organelle membrane, integral component of the organelle membrane, peroxisomal membrane, integral component of the peroxisomal membrane, intrinsic component of the peroxisomal membrane, microbody membrane, microbody part, and peroxisomal part (Figure 7a).

**Figure 7.**
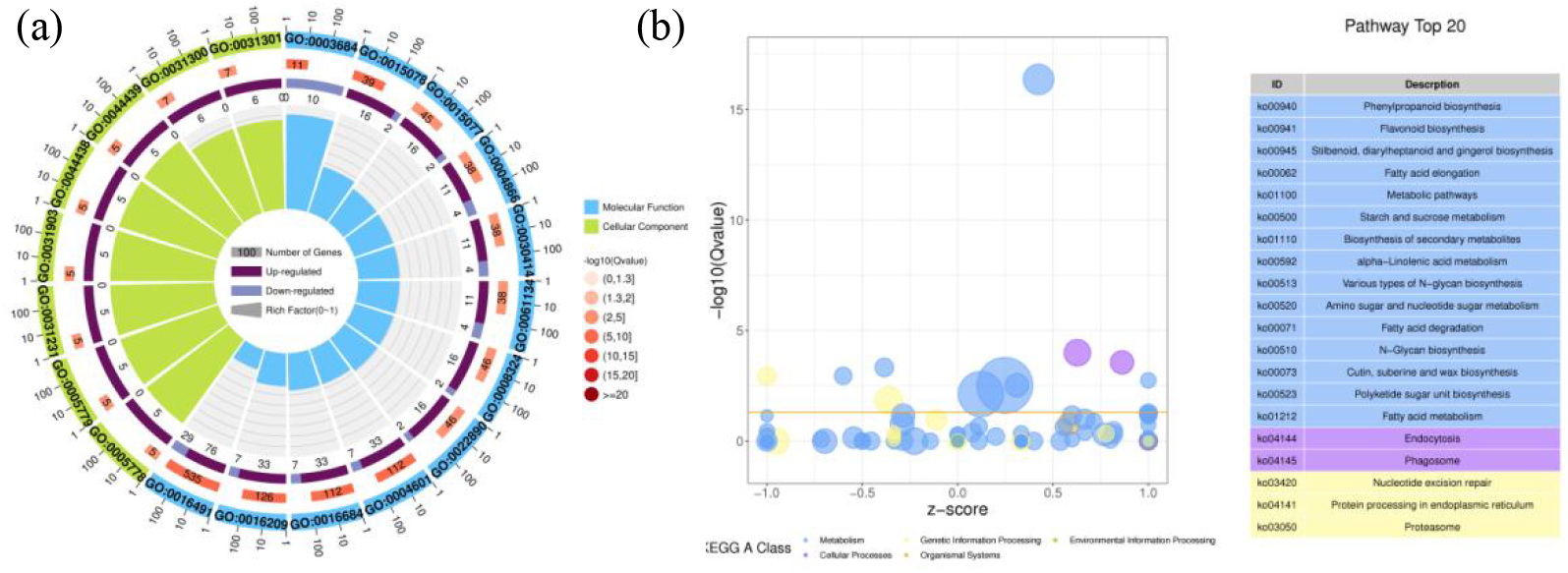
The results of GO and KEGG enrichment analyses of NS vs. FS DEPs. (a) Go enrichment circle diagram. The first circle: enrichment of the top 20 GO terms, and the outside circle is the coordinate scale of the number of genes. Different colors represent different ontologies; the second circle: the number of the GO terms in the background genes and the Q value. The more genes, the longer the bar, the smaller the Q value, and the redder the color; the third circle: bar chart of up-down gene ratio, dark purple represents the proportion of up-regulated genes, light purple represents the proportion of down-regulated genes; the specific values are shown below; the fourth circle: richfa of each GO terms rich factor (the number of differential genes in this goterm divided by all numbers), background grid lines, each grid representing 0.1. (b) Ko enrichment difference bubble diagram: (the ordinate is -log10 (Qvalue), and the abscissa is Z-score value (the ratio of the difference between the number of up-regulated genes and the number of down-regulated genes in the total difference genes). The yellow line represents the threshold value of Qvalue = 0.05. The right side is the list of the top 20 paths with Q values. Different colors represent different a classes.

KEGG annotation of DEPs from different combinations of soybean roots showed that the top 20 enriched pathways in NS vs. FS included phenylpropanoid biosynthesis, endocytosis, phagosome, flavonoid biosynthesis, fatty acid elongation, proteasome, etc. (Figure 7b). In NS vs. FS, 190 DEPs were annotated to 14 pathways (starch and sucrose metabolism, amino and nucleotide sugar metabolism, glycolysis/gluconeogenesis, pyruvate metabolism, etc.), which belong to the same carbohydrate metabolism group; 24 DEPs were annotated to 1 pathway (phytopathogen interaction), and 90 DEPs were annotated to 6 pathways (protein processing in the endoplasmic reticulum, proteasomes, protein export, etc.), which are all part of folding, sorting and degradation group; 70 DEPs were annotated to 13 pathways (fatty acid elongation, α-linolenic acid metabolism, fatty acid degradation, etc.) that are part of the lipid metabolism pathway (Table S3).

### Proteome/transcriptome association analysis

The nine quadrants are plotted according to the changes in gene expression in the transcriptome and proteome, which are divided into nine modules based on expression patterns. Each point in the graph represents a gene/protein, where genes with closely related change trends are distributed with proteins indicated by red points in quadrants 1, 3, 7 and 9, which are in agreement (quadrants 3 and 7) or opposite (quadrants 1 and 9) with significant differential expression correlations.

Genes and proteins that differed significantly during the period of highest infestation rates were integrated and analysed to construct a nine-quadrant map of multimolecular regulation and pathway interactions. In NS-vs.-FS, L-lactate dehydrogenase, laccase, polyphenol oxidase, thioredoxin, cysteine protease, and glutathione S-transferase were associated with the above four quadrants (Figure 8a). After inoculation with *F. mosseae*, GTP 3’,8-cyclase, O-acetyl serine transferase, 3’,5’-diphosphate nucleotidase, 2’,5’-diphosphate nucleotidase, Golgi PAP phosphatase, inositol polyphosphate 1 phosphatase, bifunctional oligonucleotidase and PAP phosphatase NrnA, L-3-cyanoalanine synthase, and cysteine synthase were identified. In N-vs.-NS, many pathogenesis-related proteins, chalcone-flavonoid isomerase, S-adenosylmethionine synthase, disulfide bond isomerase, and thioredoxin are associated in quadrants 1, 3, 7, and 9 (Figure 8b). These pathways contain specific regulatory information that explains the molecular mechanisms by which *F. mosseae* affects sulfur uptake in soybean.

**Figure 8.**
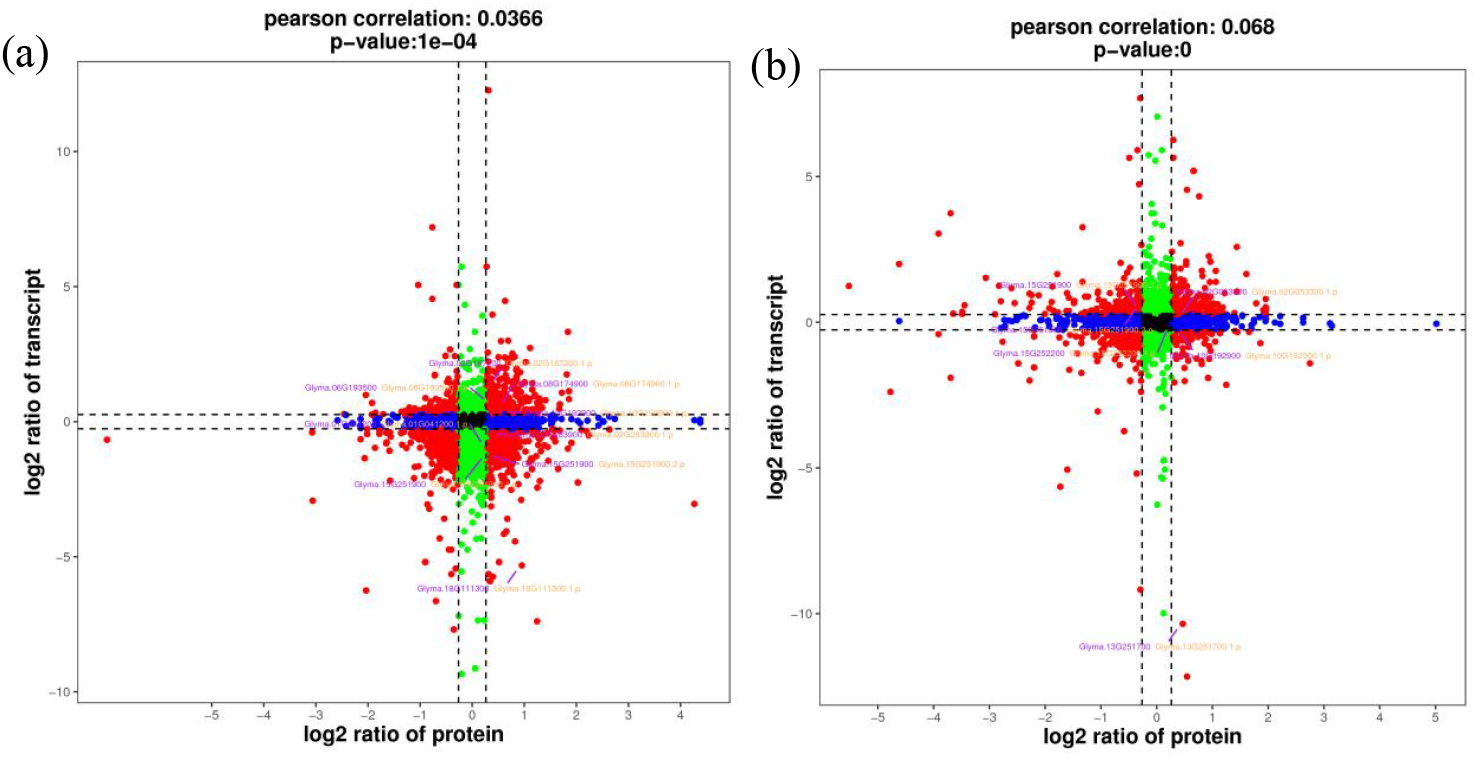
Nine-quadrant plot. (a) NS vs. FS; (b) N vs. NS. The dashed line on the abscissa represents the difference fold threshold of the transcriptome, the dashed line on the ordinate represents the difference fold threshold of the proteome, the genes / proteins outside the threshold line represent significant differences, and the genes / proteins within the threshold line represent non significant differences.

## DISCUSSION

The effect of *F. mosseae* inoculation and sulfur application on the activity of ATPS and cysteine synthase varied between the different reproductive periods. Sulfate deficiency resulted in a sharp increase in sulfate uptake and ATPS activity in soybean cells and plants and reduced the expression of seed storage proteins. After increasing sulfate, ATPS activity quickly returned to normal levels. In contrast, both moderate exogenous sulfate supply and inoculation with *F. mosseae* increased cysteine synthase activity. Thus, an additional exogenous SO_4_^2−^supply reduced the efficiency of plant sulfur assimilation, and the main way to achieve this was to reduce the activity of ATPS in the plant cytosol and chloroplasts. NS cysteine synthase activity was significantly higher after exogenous application of sulfur than when it was not applied. After inoculation with *F. mosseae*, mycorrhizae expanded the plant’s uptake area of soil nutrients, increased uptake of exogenous sulfur, contributed to sulfide accumulation, increased cysteine synthase activity, and FS cysteine synthase activity was significantly higher than that of NS.

Upon entry into plant cells, sulfate is activated to adenosyl sulfate (APS) with the release of pyrophosphate (PPi) after SO_4_^2−^ is linked to the phosphate residue on ATP via an acid-anhydride bond, catalysed by ATPS. This reaction is the only starting point for sulfate metabolism. The results of this experiment show that ATPS accumulates, and activity is enhanced in the roots of soybean plants when external sulfate is deficient. When exogenous SO_4_^2−^ is supplied, the excessive activity of ATPS in the root system is suppressed and gradually returns to normal levels. Cysteine synthase complexes are found in the mitochondria, chloroplasts and cytosol of plants. Sulfur deficiency leads to the accumulation of OAS and thus the disassembly of the cysteine complex, while increasing the supply of exogenous sulfur promotes the accumulation of sulfide and the synthesis of the enzyme complex. The results of this experiment show that sulfur supply increased the activity of cysteine synthase.

The contents of soluble protein, proline and free amino acids increased significantly (P<0.05) after *F. mosseae* inoculation, with increases of up to 1.02-3.85-fold. The results of Du et al. showed that the application of potassium sulfate (SO_4_^2−^) and sulfur powder (S^0^) was effective in increasing the height, soluble protein, dry weight and sulfur content of bok choy plants, and the enhancement effect of SO_4_^2−^ was better than that of S^0^(Du et al. 2020). This is consistent with the results of this experiment.

The sulfur-related pathways that were significantly enriched in N vs. NS and NS vs. FS were sulfur metabolism and cysteine and methionine metabolism. Subsequently, the author compared two comparable combinations of these pathways of interest.

After inoculation with AMF, both the cysteine synthase and serine O-acetyltransferase genes were upregulated in cysteine and methionine metabolism, whereas the cysteine synthase gene was downregulated after exogenous addition of thiamin. In plant-pathogen interactions, the pathogenesis-related protein 1 (PR1) gene was upregulated by AMF inoculation. In the MAPK signalling pathway - plant, inoculation with AMF activates the first line of defence for plant immunity, the receptor kinase LRR receptor-like serine (FLS2) gene, a pattern recognition receptor localized through the cytoplasmic membrane, while the PR1 gene is upregulated in expression by AMF inoculation (Table S1).

Sulfur is an essential element for plant growth and development; plants synthesize cysteine and methionine residues by reductive assimilation of sulfate, and cysteine biosynthesis is the final step in the assimilation of inorganic sulfate by plants (Bermúdez et al. 2010). In addition to binding to proteins, cysteine is the basis for the biosynthesis of many sulfur-containing molecules and cofactors (Selles et al. 2019). O-acetyl serine sulfhydrylase (OASS; O-acetylserine sulfhydrylase; EC 2.5.1.47) and acetyl serine transferase (SAT; serine Oacetyltransferase; EC 2.3.1.30) are key enzymes in the synthesis of cysteine. As the first step in the cysteine biosynthetic pathway, acetyl serine transferase catalyses the formation of O-acetyl serine from acetyl coenzyme A and serine (Jez 2019). The synthesis of cysteine involves the substitution of the O-acetyl group of O-acetyl serine with a sulfide, a reaction catalysed by pyridoxal 5’ phosphate (PLP) dependent on O-acetyl serine sulfhydrylase. The results of this experiment confirm that the pathways of cysteine and methionine metabolism and sulfur metabolism were both annotated to show significant downregulation of cysteine synthase (*cysK*) expression following the exogenous addition of thiamin. After inoculation with *F. mosseae*, O-acetyl serine transferase (*cysE*), cysteine synthase (*cysK, cysO, ATCYSC1*) and o-phosphatidylserine sulfhydrylase/cystathionine β synthase (*ATCYSC1)* were significantly upregulated in the NS vs. FS sulfur metabolism pathway, whereas 3’ (2’), 5’-diphosphate nucleotidase (*cysQ, MET22, BPNT1*), Golgi PAP phosphatase (*IMPAD1, IMPA3)* and other phosphatase genes were significantly downregulated.

In plants, autophagy plays a ‘pro-death’ and ‘pro-survival’ role in controlling programmed cell death associated with immunity triggered by microbial effectors (Lenz et al. 2011). Autophagy plays an important role in nutrient cycling during plant nitrogen or carbon starvation and in response to abiotic stresses and regulates age- and immune-related programmed cell death, which is important in plant resistance to biotrophic pathogens(Lai et al. 2011). The results of this experiment show that following inoculation with *F. mosseae*, fungal infestation triggers sensitive soybean root systems to initiate defence responses related to the pathway, including autophagic biological processes in response to other eukaryotes. In addition to autophagy, the plant MAPK signalling pathway not only responds to a variety of abiotic and biotic stresses but also transmits and amplifies external environmental signals in response(Mcneece et al. 2019). The MAPK cascade regulates growth, development, stress responses and immunity by sensing signals from upstream regulators and transmitting phosphorylated signals to downstream signalling components(Wang et al. 2020; Chang et al. 2019). Based on the MAPK signalling pathway map, we hypothesize that *F. mosseae* triggers basic plant defence responses similar to those of pathogenic infestation, the plant MAPK signalling pathway, flavonoid and flavanol biosynthesis, flavonoid biosynthesis, and tropane, piperidine and pyridine alkaloid biosynthesis.

Microtubules are an important component of the cytoskeleton and play an important role in numerous biological processes, such as polar growth, material transport, mitosis and maintenance of cell morphology in fungi (Wang. et al. 2019). The transcriptome sequencing results of this experiment showed that DEGs were significantly enriched in the cytoskeleton, microtubule cytoskeleton and cytoskeleton protein binding GO terms after inoculation with *F. mosseae*. Auxin response factor (ARF) plays a critical role in plant growth and development. ARF 6 genes are associated with the growth of the apical meristem, development of vascular tissues, release of apical shoot dormancy and development of floral organs, and ARF 2 regulates leaf senescence and floral organ abscission(Li et al. 2015; Hao et al. 2020; Guan et al. 2013). After analysis of NS vs. FS, GO results showed that DEGs were significantly enriched in the regulation of ARF protein signalling, ARF protein signalling, and ARF guanylate exchange factor activity after inoculation with *F. mosseae*.

The above results provide preliminary insights into the phenotypic differences and molecular mechanisms of soybean root response to sulfur nutrient supply and *F. mosseae* inoculation under different conditions. It was verified that *F. mosseae* could promote the uptake of soil sulfur in continuously cropped soybean at the physiological and biochemical levels and improve the biological indicators and yield quality of soybean. The effect of *F. mosseae* on the uptake, translocation and assimilation of sulfur in soybean roots was explored at the transcriptome and proteome levels, and the reasons for the differential expression of sulfur-related proteins and genes were elucidated.

## MATERIALS AND METHODS

### Plant material, AMF inoculum and experimental design

‘HN48’ plants were grown at the Sugar Research Institute experimental station of Harbin Institute of Technology (125°42’-130°10’ E, 44°04’-46°40’ N, Nangang District, Harbin City, China). The test strain *F. mosseae was* provided by the Institute of Plant Nutrition and Resources, Beijing Academy of Agriculture and Forestry Sciences, using the legume *Medicago sativa L*. The inoculum consisted of spores, mycelia and substrate (air-dried soil, sand and vermiculite). Air-dried soil of three years of continuous HN48 soybeans versus the first year of HN48 soybeans (with maize as the previous crop) was used for the pot experiment.

Each pot was filled with 2.5 kg of soil, and HN 48 soybean seeds were selected for their fullness before sowing. HN48 was sown in first-year soybean soil as a control CK; HN48 was sown in three-year-old soybean soil as N; HN48 was sown in three-year-old soybean soil with 0.5 mmol/L MgSO_4_ as NS; HN48 was sown in three-year-old soy*bean soil* inoculated with *F. mosseae* and 0.5 mmol/L MgSO_4_ as FS. CK and N were supplemented with equal amounts of 0.5 mmol/L MgCl_2_; CK, N and NS were inoculated with equal amounts of inactivated *F. mosseae*. The collected samples were washed with deionized water to remove surface dust and stored at -80 °C in a refrigerator after liquid nitrogen treatment.

### Plant sampling and parameter determination

Soybean root samples were taken weekly from 30 d after sowing until 90 d. *F. mosseae* infestation was determined using the acidic magenta staining method. Samples were taken every 15 d after 30 d of sowing, and the 2,3,5-triphenyl tetrazolium chloride (TTC) staining method was used to assess root activity(Bai et al.1994). The total sulfate content of soybean leaves and roots was determined using the BaSO_4_ turbidimetric method. The soluble protein content of soybean leaves and roots was determined by Kormas Brilliant Blue colorimetry (Bradford, 1976). ATP sulfurylase (ATPS) activity was determined by Lappartient’s method(Lappartient et al.1996). Cysteine synthase (CSase) activity was determined by Warrilow’s method(Warrilow et al.1998)..

### Construction of libraries, RNA-seq and identification of DEGs

The methods of construction of libraries and RNA-sequencing are provided in Methods S1. We deposited our sequencing data uploaded to the NCBI Sequence Read Archive (SRA) under accession number SRP312573. RNA differential expression analysis was performed by DESeq2(Love et al. 2014) software between two different groups (and by edgeR(Robinson et al. 2010) between two samples). Genes/transcripts with the parameter of false discovery rate (FDR) below 0.05 and absolute fold change≥2 were considered DEGs/transcripts.

### Verification of RNA-seq data by qPCR

To verify the reliability of the RNA-seq data, qRT-PCR was performed to validate gene expression. This experiment used TransZol Up (TransGen Biotech, Beijing) to extract total RNA from soybean roots. RNA was reverse transcribed with the aid of *EasyScript®* All-in-One First-Strand cDNA Synthesis SuperMix for qPCR (TransGen Biotech, Beijing, China), and the reaction system is described in Table 1. Heat for 5 s at 42°C to inactivate EasyScript® RT/RI and gDNA Remover.

**Table 1.** Composition of the RNA reverse transcription reaction system in soybean roots.

| Reagents | Volume (μL) |
| --- | --- |
| Total RNA/mRNA | 1 μg |
| 5 × EasyScript® All-in-One SuperMix for qPCR | 4 μL |
| gDNA Remover | 1 μL |
| RNase-free Water | Up to 20 $\mu$ L |

### Protein extraction and digestion

Total proteins were extracted using the cold acetone method. Samples were ground to powder in liquid nitrogen and then dissolved in 2 mL lysis buffer (8 M urea, 2% SDS, 1x Protease Inhibitor Cocktail (Roche Ltd. Basel, Switzerland), followed by sonication on ice for 30 min and centrifugation at 13 000 rpm for 30 min at 4 °C. The supernatant was transferred to a fresh tube. For each sample, proteins were precipitated with ice-cold acetone at -20 °C overnight. The precipitates were cleaned with acetone three times and redissolved in 8 M urea by sonication on ice. Protein quality was examined with SDS-PAGE. A Protein Assay Kit was used to determine the protein concentration of the supernatant. Fifty micrograms of protein extracted from cells was suspended in 50 μL of solution. Then, the sample was precipitated using 300 μL of pre-preparation solution. The precipitate was washed twice with cold acetone and then resuspended in 50 mM. Finally, the proteins were digested with sequence-grade modified trypsin (Promega, Madison, WI) at a substrate/enzyme ratio of 50:1 (w/w) at 37 °C for 16 h.

### LC-MS/MS, data and functional analysis

The peptides were redissolved in 30 μL solvent A (A: 0.1% formic acid in water) and analysed by online nanospray LC-MS/MS on an Orbitrap Fusion Lumos coupled to an EASY-nLC 1200 system (Thermo Fisher Scientific, MA, USA). A 3 μL peptide sample was loaded onto the analytical column (Acclaim PepMap C18, 75 μm × 25 cm) with a 120-min gradient, from 5% to 35% B (B: 0.1% formic acid in ACN). The column flow rate was maintained at 200 nL/min with a column temperature of 40 °C. The electrospray voltage was 2 kV versus the inlet of the mass. The mass spectrometer was run under data-independent acquisition mode and automatically switched between MS and MS/MS mode.

Raw DIA data were processed and analysed by Spectronaut X (Biognosys AG, Switzerland) with default parameters. Data extraction was determined by Spectronaut X based on extensive mass calibration. The Q value (FDR) cut-off at the precursor and protein levels was 1%. Decoy generation was set to mutate, which is similar to scrambled but will only apply a random number of AA position swamps (min=2, max=length/2). The average top 3 filtered peptides that passed the 1% Q value cut-off were used to calculate the major group quantities. After the t test, DEPs were filtered if their Q value <0.05 and absolute AVG log2 ratio 0.58 Proteins were annotated against GO and KEGG databases to obtain their functions. Significant GO functions and pathways were examined within DEPs with q value≤0.05.

## Data availability

The RNA-seq data obtained in the present study were deposited in the NCBI Sequence Read Archive (SRA, https://www.ncbi.nlm.nih.gov/sra/) under the accession number SRP312573.

## Supplemental Data

**Supplemental Figure S1**. Physiological indices of soybean plants. (a) Plant height; (b) plant dry weight; (c) plant fresh weight.

**Supplemental Figure S2**. Content of soluble protein in soybean plants. (a) Leaf soluble protein; (b) root soluble protein.

**Supplemental Figure S3**. Trend analysis of gene expression.

**Supplemental Figure S4**. Relative expression levels of five target genes determined by qRT-PCR: (a) LBA; (b) CHI4A; (c) HSP; (d) GSTU9; (e) SAMS.

**Supplemental Table S1**. The results of NS vs. FS differentially expressed gene GO enrichment analysis.

**Supplemental Table S2**. The result of NS vs. FS differentially expressed gene KEGG enrichment analysis.

**Supplemental Table S3**. The results of NS vs. FS differentially expressed gene GSEA-GO analysis.

**Supplemental Table S4**. The results of NS vs. FS differentially expressed gene GSEA-KEGG analysis.

**Supplemental Methods S1**. The methods of construction of libraries and RNA-sequencing.

## ACKNOWLEDGMENTS

Authors appreciated the Institute of Plant Nutrition and Resources, Beijing Academy of Agriculture and Forestry Sciences for providing fungal strains.

## AUTHOR CONTRIBUTION

BY Cai and XQ Zhang planned and designed the research. XQ Zhang, CC Lu, RL Liu and ZX Sun performed research. BY Cai and XQ Zhang conducted fieldwork, analysed data etc. XQ Zhang wrote the manuscript.

